# Genomic diversity of *Escherichia coli* isolates from healthy children in rural Gambia

**DOI:** 10.1101/2020.08.28.271627

**Authors:** Ebenezer Foster-Nyarko, Nabil-Fareed Alikhan, Usman Nurudeen Ikumapayi, Sarwar Golam, M Jahangir Hossain, Catherine Okoi, Peggy-Estelle Tientcheu, Marianne Defernez, Justin O’Grady, Martin Antonio, Mark J. Pallen

## Abstract

Little is known about the genomic diversity of *Escherichia coli* in healthy children from sub-Saharan Africa, even though this is pertinent to understanding bacterial evolution and ecology and their role in infection. We isolated and whole-genome sequenced up to five colonies of faecal *E. coli* from 66 asymptomatic children aged three-to-five years in rural Gambia (n=88 isolates from 21 positive stools). We identified 56 genotypes, with an average of 2.7 genotypes per host. These were spread over 37 seven-allele sequence types and the *E. coli* phylogroups A, B1, B2, C, D, E, F and *Escherichia* cryptic clade I. Immigration events accounted for three-quarters of the diversity within our study population, while one-quarter of variants appeared to have arisen from within-host evolution. Several study strains were closely related to isolates that caused disease in humans or originated from livestock. Our results suggest that within-host evolution plays a minor role in the generation of diversity than independent immigration and the establishment of strains among our study population. Also, this study adds significantly to the number of commensal *E. coli* genomes, a group that has been traditionally underrepresented in the sequencing of this species.

## Introduction

Ease of culture and genetic tractability account for the unparalleled status of *Escherichia coli* as “the biological rock star”, driving advances in biotechnology (1), while also providing critical insights into biology and evolution (2). However, *E. coli* is also a widespread commensal, as well as a versatile pathogen, linked to diarrhoea (particularly in the under-fives), urinary tract infection, neonatal sepsis, bacteraemia and multi-drug resistant infection in hospitals (3-5). Yet, most of what we know about *E. coli* stems from the investigation of laboratory strains, which fail to capture the ecology and evolution of this key organism “in the wild” (6). What is more, most studies of non-lab strains have focused on pathogenic strains or have been hampered by low-resolution PCR methods, so we have relatively few genomic sequences from commensal isolates, particularly from low-to middle-income countries (7-13).

We have a broad understanding of the population structure of *E. coli*, with eight significant phylogroups loosely linked to ecological niche and pathogenic potential (B2, D and F linked to extraintestinal infection; A and B1 linked to severe intestinal infections such as haemolytic-uraemic syndrome) (14-17). All phylogroups can colonise the human gut, but it remains unclear how far commensals and pathogenic strains compete or collaborate—or engage in horizontal gene transfer—within this important niche (18, 19).

Although clinical microbiology typically relies on single-colony picks (which has the potential to underestimate species diversity and transmission events), within-host diversity of *E. coli* in the gut is crucial to our understanding of inter-strain competition and co-operation and also for accurate diagnosis and epidemiological analyses. Pioneering efforts using serotyping and molecular typing have shown that normal individuals typically harbour more than one strain of *E. coli* (20-22), with one individual carrying 24 distinct clones (22-24). More recently, whole-genome sequencing has illuminated molecular epidemiological investigations (9), adaptation during and after infection (25, 26), as well as the intra-clonal diversity in healthy hosts (27).

There are two plausible sources of within-host genomic diversity. Although a predominant strain usually colonises the host for extended periods (28), successful immigration events mean that incoming strains can replace the dominant strain or co-exist alongside it as minority populations (29). Strains originating from serial immigration events are likely to differ by hundreds or thousands of single-nucleotide polymorphisms (SNPs). Alternatively, within-host evolution can generate clouds of intra-clonal diversity, where genotypes differ by just a handful of SNPs (20).

Most relevant studies have been limited to Western countries, except for a recent report from Tanzania (21), so little is known about the genomic diversity of *E. coli* in sub-Saharan Africa. The Global Enteric Multicenter Study (GEMS) (30, 31) has documented a high burden of diarrhoea attributable to *E. coli* (including *Shigell*a) among children from the Gambia, probably as a result of increased exposure to this organism through poor hygiene and frequent contact with animals and the environment. In also facilitating access to stool samples from healthy Gambian children, the GEMS study has given us a unique opportunity to study within-host genomic diversity of commensal *E. coli* in this setting.

## Methods

### Study population

We initially selected 76 faecal samples from three-to five-year-old asymptomatic Gambian children, who had been recruited from Basse, Upper River Region, the Gambia, into the GEMS study (30) as healthy controls from December 1, 2007, to March 3, 2011. Samples had been collected according to a previously described sampling protocol (32). Archived stool samples were retrieved from -80°C storage and allowed to thaw on ice. A 100-200 mg aliquot from each sample was transferred aseptically into 1.8ml Nunc tubes for microbiological processing below (Figure 1). Ten of the original 76 samples proved unavailable for processing in this study.

**Figure 1.**
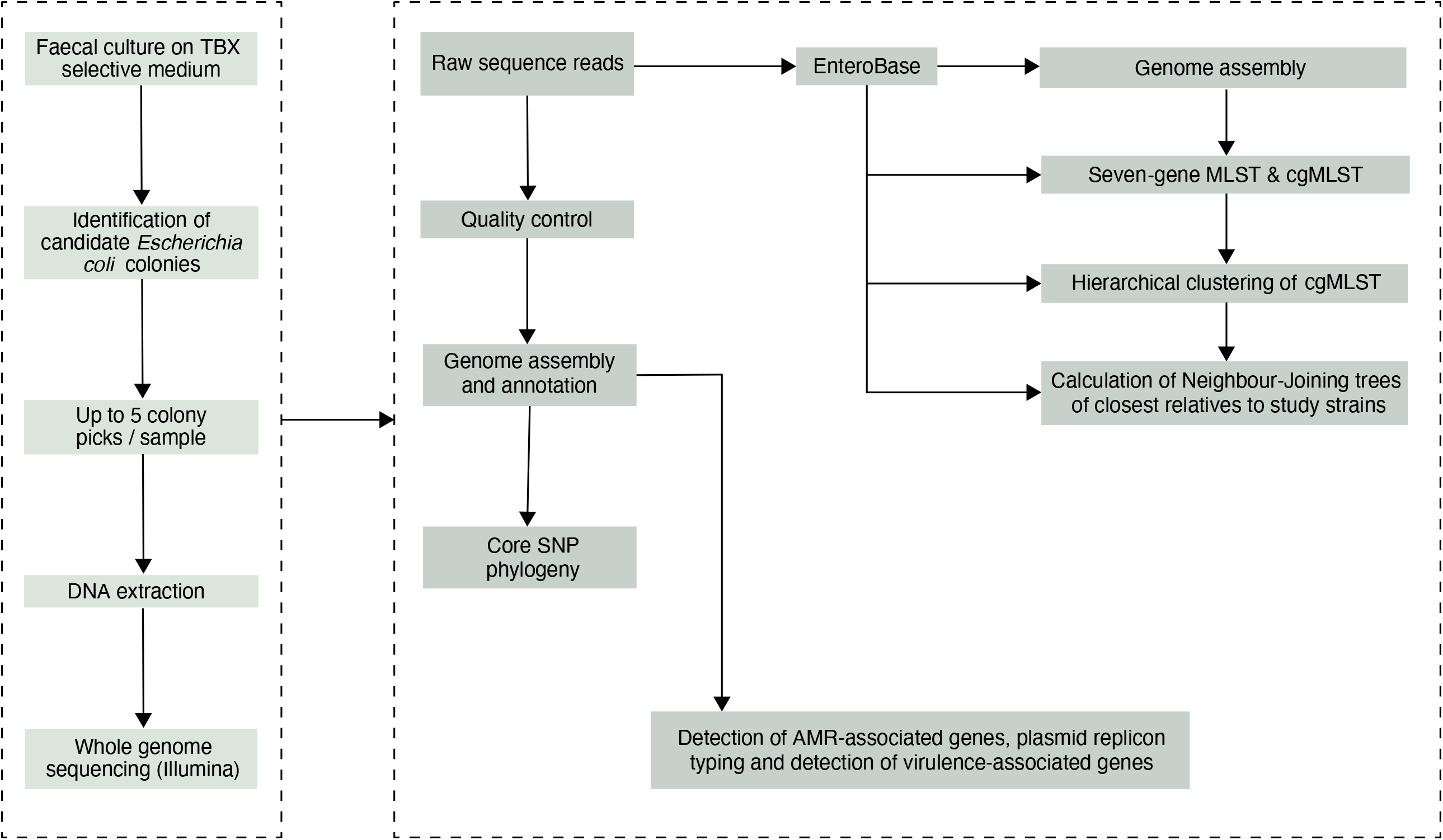
The study sample processing flow diagram.

### Bacterial growth and isolation

1 ml of physiological saline (0.85%) was added to each sample tube and vigorously vortexed at 4200 rpm for at least 2 minutes. Next, the homogenised sample suspensions were taken through four ten-fold dilution series. A100 µl aliquot from each dilution was then spread evenly on a plate of tryptone-bile-X-glucuronide differential and selective agar. The inoculated plates were incubated overnight at 37°C under aerobic conditions. Colony counts were performed on the overnight cultures for each serial dilution for translucent colonies with entire margins and blue-green pigmentation indicative of *E. coli*. Up to five representative colonies were selected from each sample and sub-cultured on MacConkey agar overnight at 37°C before storing in 20% glycerol broth at -80°C. Individual isolates were assigned a designation comprised of the subject ID followed by the colony number (“1-5”).

### Genomic DNA extraction and genome sequencing

Broth cultures were prepared from pure, fresh cultures of each colony-pick in 1 ml Luria-Bertani broth and incubated overnight to attain between 10^9^ – 10^10^ cfu per ml. Genomic DNA was then extracted from the overnight broth cultures using the lysate method described in (33). The eluted DNA was quantified by the Qubit high sensitivity DNA assay kit (Invitrogen, MA, USA) and sequenced on the Illumina NextSeq 500 instrument (Illumina, San Diego, CA) as described previously (34).

Following Dixit et al. (20), we sequenced a random selection of isolates twice, using DNA obtained from independent cultures, to help in the determination of clones and the analysis of within-host variants (Supplementary File 5). Bioinformatic analyses of the genome sequences were carried out on the Cloud Infrastructure for Microbial Bioinformatics (CLIMB) platform (35).

### Phylogenetic analysis

The paired 150bp reads were quality checked and assembled, as previously described (34). Snippy v4.3.2 (https://github.com/tseemann/snippy) was used for variant calling, using the complete genome sequence of commensal *E. coli* str. K12 substr. MG1655 as a reference strain (NCBI accession: NC_000913.3) and to generate a core-genome alignment, from which a maximum-likelihood phylogeny with 1000 bootstrap replicates was reconstructed using RAxML v8.2.4 (36), based on a general time-reversible nucleotide substitution model. The phylogenetic tree was rooted using the genomic sequence of *E. fergusonii* as an outgroup (NCBI accession: GCA_000026225.1). The phylogenetic tree was visualised in FigTree v1.4.3 (https://github.com/rambaut/figtree/) and annotated in RStudio v3.5.1 and Adobe Illustrator v 23.0.3 (Adobe Inc., San Jose, California). For visualisation, a single colony was chosen to represent replicate colonies of the same strain (ST) with identical virulence, plasmid and antimicrobial resistance profiles and a de-replicated phylogenetic tree reconstructed using the representative isolates.

### Multi-locus sequence typing, Clermont typing and SNPs

The merged reads were uploaded to EnteroBase (37), where *de novo* assembly and genome annotation were carried out, and *in-silico* multi-locus sequence types (STs) assigned based on the Achtman scheme, allocating new sequence types (ST) if necessary. EnteroBase assigns phylogroups using ClermontTyper and EzClermont (38, 39) and unique core-genome MLST types based on 2, 513 core loci in *E. coli*. Publicly available *E. coli* sequences in EnteroBase (http://enterobase.warwick.ac.uk/species/index/ecoli) (37) were included for comparative analysis, including 23 previously sequenced isolates obtained from diarrhoeal cases recruited in the GEMS study in the Gambia (Supplementary File 1).

We computed pairwise single nucleotide polymorphism (SNP) distances between genomes from the core-genome alignment using snp-dists v0.6 (https://github.com/tseemann/snp-dists). For the duplicate sequence reads of the same strains, we used SPAdes v3.13.2 (40) to assemble each set of reads and map the raw sequences from one sequencing run to the assembly of the other run and vice versa, as described previously (20). SNPs were detected using the CSIPhylogeny tool (https://cge.cbs.dtu.dk/services/CSIPhylogeny/) and compared between the two steps, counting only those SNPs that were detected in both sets of reads as accurate.

### Accessory gene content

We used ABRicate v0.9.8 (https://github.com/tseemann/abricate) to predict virulence factors, acquired antimicrobial resistance (AMR) genes and plasmid replicons by scanning the contigs against the VFDB, ResFinder and PlasmidFinder databases respectively, using an identity threshold of ≥ 90% and a coverage of ≥ 70%. Virulence factors and AMR genes were plotted next to the phylogenetic tree using the ggtree, ggplot2 and phangorn packages in RStudio v3.5.1. We calculated co-occurrence of AMR genes among study isolates and visualised this as a heat map using RStudio v 3.5.1.

### Population structure and comparison of commensal and pathogenic strains

We assessed the population structure using the hierarchical clustering algorithm in EnteroBase. Briefly, the isolates were assigned stable population clusters at eleven levels (from HC0 to HC 2350) based on pairwise cgMLST allelic differences. Hierarchical clustering at 1100 alleles differences (HC1100) resolves populations into cgST complexes, the equivalent of clonal complexes achieved with the legacy MLST clustering approaches (37). We reconstructed neighbour-joining phylogenetic trees using NINJA (41), based on clustering at HC1100 to display the population sub-clusters at this level as an indicator of the genomic diversity within our study population and to infer the evolutionary relationship among our strains and others in the public domain.

Next, we interrogated the HC1100 clusters that included both pathogenic and commensal *E. coli* strains recovered from the GEMS study. For the clusters that encompassed commensal and pathogenic strains belonging to the same ST, we reconstructed both neighbour-joining and SNP phylogenetic trees to display the genetic relationships among these strains. We visualised the accessory genomes for the overlapping STs mentioned above to determine genes associated with phages, virulence factors and AMR. The resulting phylogenetic trees were annotated in Adobe Illustrator v 23.0.3 (Adobe Inc., San Jose, California).

### Ethical statement

The study was approved by the joint Medical Research Council Unit The Gambia-Gambian Government ethical review board.

## Results

### Population structure

The study population included 27 females and 39 males (Table 1). All but one reported the presence of a domestic animal within the household. Twenty-one samples proved positive for the growth of *E. coli*, yielding 88 isolates. We detected 37 seven-allele sequence types (STs) among the isolates, with a fairly even distribution (Figure 2). Five STs were completely novel (ST9274, ST9277, ST9278, ST9279 and ST9281). These study strains were scattered over all the eight main phylogroups of *E. coli* (Table 2). Hierarchical clustering of core genomic STs revealed twenty-seven cgST clonal complexes (Supplementary File 2).

**Table 1:** Characteristics of the study population

| Sample ID | Lab ID | Age (months) | Gender | Bristol stool index | Domestic animal within household | Enrolment date |
| --- | --- | --- | --- | --- | --- | --- |
| 102135 | H1 | 43 | Female | Thick liquid | Goat, sheep | 18-Feb-09 |
| 102650 | H2 | 45 | Female | Soft | Goat, sheep, donkey | 27-Jul-09 |
| 103296 | H3 | 44 | Male | Soft | Goat, horse, donkey, rodent | 27-Apr-10 |
| 103298 | H4 | 44 | Male | Formed | Sheep, fowl, horse, donkey, rodent | 27-Apr-10 |
| 103621 | H5 | 37 | Female | Soft | Sheep, fowl, rodent | 01-Sep-10 |
| 103650 | H6 | 48 | Female | Soft | Fowl, donkey, rodent | 29-Sep-10 |
| 103649 | H7 | 45 | Female | Soft | Goat, sheep, fowl, horse, rodent | 29-Sep-10 |
| 103071 | H8 | 53 | Male | Formed | Goat, sheep, fowl | 15-Jan-10 |
| 103622 | H9 | 39 | Female | Soft | Goat, sheep | 01-Sep-10 |
| 100167 | H10 | 40 | Female | Soft | Goat, sheep, fowl | 01-Feb-08 |
| 100217 | H11 | 57 | Male | Formed | Cat, fowl, horse, rodent | 21-Feb-08 |
| 100230 | H12 | 51 | Male | Soft | Goat, sheep, cat, fowl, rodent | 28-Feb-08 |
| 100612 | H13 | 55 | Female | Formed | Goat, sheep, dog, fowl, horse, donkey, rodent | 16-Aug-08 |
| 100162 | H14 | 47 | Female | Thick liquid | Sheep, horse, donkey, rodent | 30-Jan-08 |
| 102255 | H15 | 42 | Male | Formed | Goat, sheep, fowl, horse, donkey, rodent | 26-Mar-09 |
| 102250 | H16 | 39 | Male | Formed | Fowl | 25-Mar-09 |
| 102114 | H17 | 54 | Male | Formed | Rodent | 12-Feb-09 |
| 102123 | H18 | 37 | Female | Soft | Goat, sheep, fowl, rodent | 14-Feb-09 |
| 103282 | H19 | 43 | Male | Formed | Goat, sheep, dog, cat, cow, fowl, | 22-Apr-10 |
| 100817 | H20 | 44 | Male | Soft | Dog, fowl | 03-Dec-08 |
| 100816 | H21 | 40 | Male | Soft | Goat, sheep, cow, fowl, horse, donkey, rodent | 03-Dec-08 |
| 102836 | H22 | 47 | Male | Thick liquid | Fowl, rodent | 12-Oct-09 |
| 102837 | H23 | 41 | Male | Thick liquid | Sheep, fowl, rodent | 12-Oct-09 |
| 102843 | H24 | 44 | Male | Soft | Fowl, rodent | 13-Oct-09 |
| 102907 | H25 | 36 | Male | Soft | Goat, sheep, fowl | 05-Nov-09 |
| 102905 | H26 | 37 | Male | Soft | Goat, sheep, fowl | 05-Nov-09 |
| 102262 | H27 | 38 | Male | Formed | Goat, sheep, rodent | 01-Apr-09 |
| 102728 | H28 | 41 | Male | Soft | Goat, fowl | 24-Aug-09 |
| 102729 | H29 | 41 | Male | Soft | Goat, dog, cat, fowl, donkey | 24-Aug-09 |
| 100806 | H30 | 55 | Male | Soft | Goat, sheep, dog, fowl | 21-Nov-08 |
| 102053 | H31 | 37 | Female | Formed | Cow, fowl, donkey, rodent | 29-Jan-09 |
| 102052 | H32 | 38 | Female | Formed | Goat, sheep, cow, fowl, donkey, rodent | 29-Jan-09 |
| 102511 | H33 | 37 | Male | Soft | Fowl, horse, donkey, rodent | 19-Jun-09 |
| 102649 | H34 | 37 | Male | Soft | Fowl, horse, donkey, rodent | 27-Jul-09 |
| 102454 | H35 | 52 | Male | Soft | Sheep, fowl, donkey, rodent | 02-Jun-09 |
| 102459 | H36 | 51 | Male | Formed | Goat, sheep, dog, cat, cow, horse, donkey, rodent | 04-Jun-09 |
| 100303 | H37 | 58 | Male | Formed | Sheep, fowl | 08-Apr-08 |
| 100320 | H38 | 42 | Female | Formed | Sheep, fowl, rodent | 19-Apr-08 |
| 100319 | H39 | 45 | Female | Formed | Goat, sheep, fowl, rodent | 17-Apr-08 |
| 103081 | H40 | 39 | Female | Thick liquid | Goat, sheep, fowl, horse, donkey, rodent | 20-Jan-10 |
| 103082 | H41 | 39 | Female | Thick liquid | Goat, sheep, fowl, horse, donkey, rodent | 20-Jan-10 |
| 100663 | H42 | 36 | Male | Thick liquid | Goat, sheep, fowl, donkey | 10-Sep-08 |
| 100072 | H43 | 51 | Female | Formed | Goat, cow, fowl, rodent | 03-Jan-08 |
| 103171 | H44 | 36 | Female | Soft | Goat, sheep, rodent, fowl, rodent | 18-Feb-10 |
| 103172 | H45 | 36 | Female | Soft | Goat, sheep, fowl, rodent | 18-Feb-10 |
| 103292 | H46 | 39 | Male | Soft | Goat, sheep, fowl | 23-Apr-10 |
| 102952 | H47 | 36 | Male | Soft | Goat, sheep, fowl, rodent | 20-Nov-09 |
| 102953 | H48 | 37 | Male | Soft | Goat, sheep, fowl, rodent | 20-Nov-09 |
| 102964 | H49 | 40 | Female | Formed | Goat, fowl, rodent | 26-Nov-09 |
| 102966 | H50 | 37 | Female | Formed | Goat, sheep, fowl, horse, donkey, rodent | 22-Apr-10 |
| 103281 | H51 | 44 | Male | Formed | Goat, sheep, dog, cat, fowl | 22-Apr-10 |
| 100540 | H52 | 43 | Male | Soft | Goat, sheep, fowl, rodent | 22-Jul-08 |
| 103123 | H53 | 38 | Male | Soft | Sheep | 03-Feb-10 |
| 103124 | H54 | 36 | Male | Soft | Fowl | 03-Feb-10 |
| 102089 | H55 | 38 | Female | Soft | Goat, cow, fowl, horse, donkey, rodent | 05-Feb-09 |
| 103297 | H56 | 38 | Male | Soft | Goat, sheep, fowl, horse, donkey, rodent | 27-Apr-10 |
| 102251 | H57 | 39 | Male | Formed | Fowl | 25-Mar-09 |
| 103602 | H58 | 38 | Female | Formed | Goat, sheep, cow, fowl | 26-Aug-10 |
| 103600 | H59 | 39 | Female | Formed | Goat, sheep, fowl | 26-Aug-10 |
| 100026 | H60 | 49 | Female | Soft | Goat, sheep, cow, fowl | 14-Dec-07 |
| 102102 | H61 | 47 | Female | Opaque watery | None | 11-Feb-09 |
| 102263 | H62 | 38 | Male | Formed | Horse, donkey, rodent | 01-Apr-09 |
| 103070 | H63 | 58 | Male | Soft | Goat, sheep, fowl | 15-Jan-10 |
| 103130 | H64 | 40 | Male | Soft | Sheep, fowl | 03-Feb-10 |
| 102051 | H65 | 36 | Female | Formed | Goat, sheep, dog, cat, cow, fowl, donkey, rodent | 29-Jan-09 |
| 102524 | H66 | 36 | Male | Soft | Goat, sheep, fowl, horse, donkey, rodent | 24-Jun-09 |

**Table 2:** Phylogroup and sequence types of the distinct clones isolated in each patient

| *Host* | Genotype number |  |  |  |  | Number of distinct genotypes (clones) | Migration events | Within-host evolution events |
| --- | --- | --- | --- | --- | --- | --- | --- | --- |
|  | 1 | 2 | 3 | 4 | 5 |  |  |  |
| *H-2* | A (9274) | A (9274) | A (9274) | A (9274) | A (9274) | 1 | 1 | 0 |
| *H-9* | A (2705) | A (2705) | A (2705) | D (2914) | B1 (29) | 3 | 3 | 0 |
| *H-15* | B2 (9277) | B2 (9277) | B2 (9277) | Clade I (747) | Clade I (747) | 3 | 2 | 1 |
| *H-18* | D (38) | D (38) | B1 (9281) | A (9274) |  | 4 | 3 | 1 |
| *H-21* | B1 (58) | B1 (58) | B1 (223) | A (540) | D (1204) | 4 | 4 | 0 |
| *H-22* | B1 (316) | B1 (316) | B1 (316) | B1 (316) |  | 2 | 1 | 1 |
| *H-25* | A (181) | A (181) | A (181) | A (181) | B1 (337) | 4 | 2 | 2 |
| *H-26* | B1 (641) | B1 (2741) | A (10) | A (398) |  | 4 | 4 | 0 |
| *H-28* | B1 (469) | B1 (469) | B1 (469) | B1 (469) |  | 2 | 1 | 1 |
| *H-32* | B1 (101) | B1 (101) | B1 (101) | B1 (2175) | A (10) | 3 | 3 | 0 |
| *H-34* | B1 (603) | B1 (603) | B1 (603) | B1 (1727) | A (10) | 4 | 3 | 1 |
| *H-35* | A (226) |  |  |  |  | 1 | 1 | 0 |
| *H-36* | F (59) | F (59) | F (59) | F (59) | E (9278) | 3 | 2 | 1 |
| *H-37* | D (5148) | D (5148) | D (5148) | D (5148) | D (5148) | 3 | 1 | 2 |
| *H-38* | D (394) | D (394) | D (394) | D (394) | B1 (58) | 4 | 2 | 2 |
| *H-39* | B2 (452) | B2 (452) | B2 (452) | B2 (452) | B2 (452) | 2 | 1 | 1 |
| *H-40* | B1 (155) |  |  |  |  | 1 | 1 | 0 |
| *H-41* | A (43) | A (43) | A (43) | A (43) | B1 (9283) | 2 | 2 | 0 |
| *H-48* | Clade I (485) | Clade I (485) | Clade I (485) | Clade I (485) |  | 1 | 1 | 0 |
| *H-50* | C (410) | C (410) | C (410) | C (410) | B1 (515) | 2 | 2 | 0 |
| *H-55* | A (9279) |  |  |  |  | 1 | 1 | 0 |

**Figure 2.**
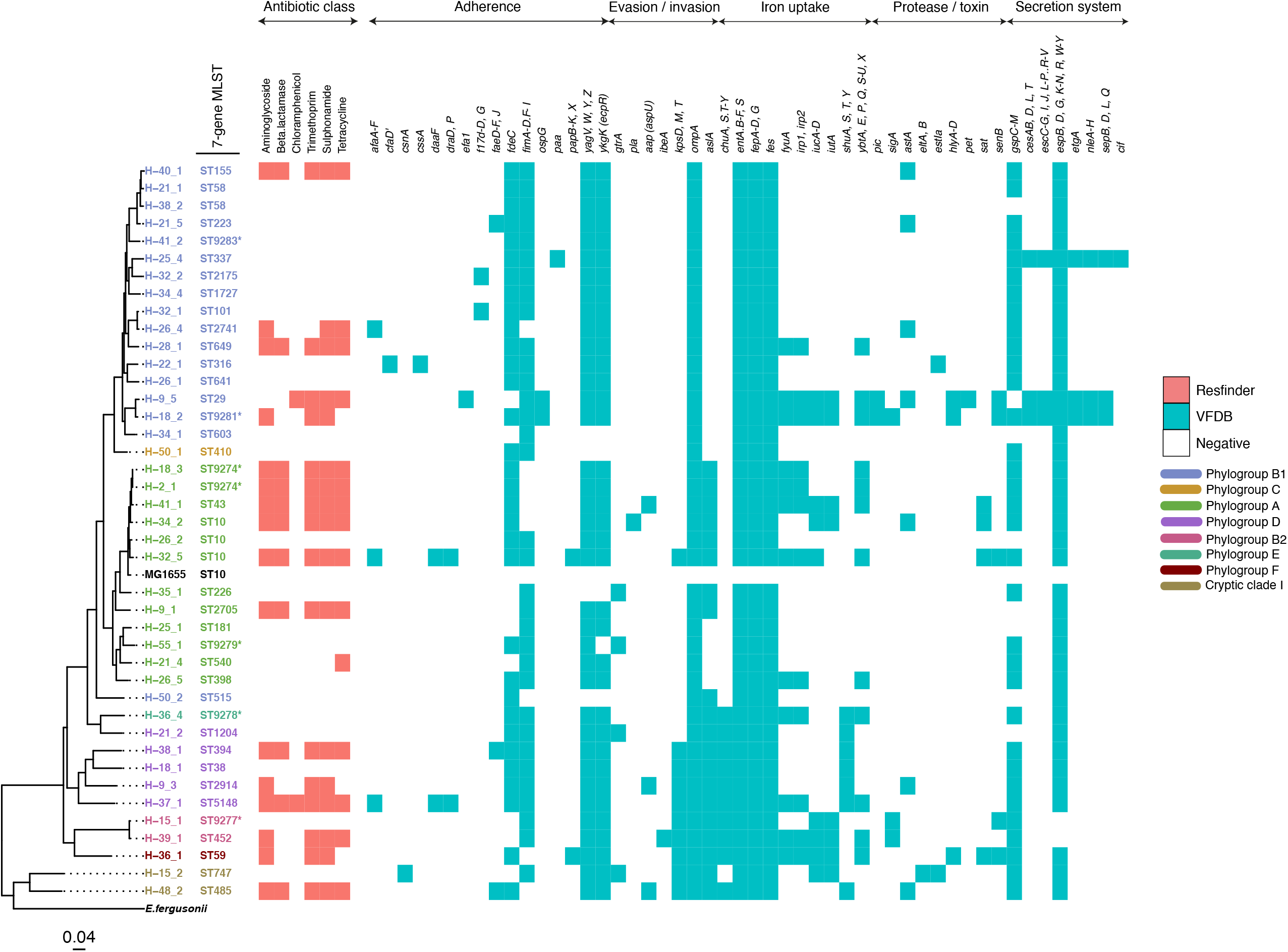
A maximum-likelihood tree depicting the phylogenetic relationships among the study isolates. The tree was reconstructed with RAxML, using a general time-reversible nucleotide substitution model and 1,000 bootstrap replicates. The genome assembly of *E. coli* str. K12 substr. MG1655 was used s as the reference, and the tree rooted using the genomic assembly of *E. fergusonii* as an outgroup. The sample names are indicated at the tip, with the respective Achtman sequence types (ST) indicated beside the sample names. The respective phylogroups the isolates belong to are indicated with colour codes as displayed in the legend. *E. coli* reference genome is denoted in black. Asterisks (*) are used to indicate novel STs. The predicted antimicrobial resistance genes and putative virulence factors for each isolate are displayed next to the tree, with the virulence genes clustered according to their function. Multiple copies of the same strain (ST) isolated from a single host are not shown. Instead, we have shown only one representative isolate from each strain. Virulence and resistance factors were not detected in the reference strain either. A summary of the identified virulence factors and their known functions are provided in Supplementary File 3.

### Within-host diversity

Just a single ST colonised nine individuals, six carried two STs, four carried four STs, and two carried six STs. We found 56 distinct genotypes, which equates to an average of 2.7 genotypes per host. Two individuals (H-18 and H-2) shared an identical strain belonging to ST9274 (zero SNP difference) (Supplementary File 4, yellow highlight), suggesting recent transfer from one child to another or recent acquisition from a common source.

We observed thirteen cases where a single host harboured two or more variants within the same SNP cloud (Table 2). Such within-host evolution accounted for around a quarter of the observed variation, with immigration explaining the remaining three quarters. 22% of within-host mutations represented synonymous changes. 43% were non-synonymous mutations, while 31% occurred in non-coding regions, and 4% represented stop-gained mutations (Supplementary File 6). The average number of SNPs among variants within such SNP clouds was 5 (range 0-18) (Table 3). However, in two subjects (H36 and H37), pairwise distances between genomes from the same ST (ST59 and ST5148) were as large as 14 and 18 SNPs respectively (Supplementary File 4, grey highlight).

**Table 3:** Pairwise SNP distances between variants arising from within-host evolution

| *Host* | Sequence type (ST) | Colonies per ST | Pairwise SNP distances between multiple colonies of the same ST |
| --- | --- | --- | --- |
| *H2* | 9274 | 5 | 0-9 |
| *H9* | 2705 | 3 | 0-1 |
| *H15* | 9277 | 3 | 0-1 |
| *H15* | 747 | 2 | 3 |
| *H18* | 38 | 2 | 3 |
| *H21* | 58 | 2 | 0 |
| *H22* | 316 | 4 | 0-3 |
| *H25* | 181 | 4 | 1-5 |
| *H28* | 469 | 4 | 0-3 |
| *H32* | 101 | 3 | 1-9 |
| *H34* | 603 | 3 | 2-8 |
| *H36* | 59 | 4 | 0-14 |
| *H37* | 5148 | 5 | 2-18 |
| *H38* | 394 | 4 | 1-3 |
| *H39* | 452 | 5 | 0-2 |
| *H41* | 43 | 4 | 0-1 |
| *H48* | 485 | 4 | 1-9 |
| *H50* | 410 | 4 | 0 |

### Accessory gene content and relationships with other strains

A quarter of our isolates were most closely related to commensal strains from humans, with smaller numbers most closely related to human pathogenic strains or strains from livestock, poultry or the environment (Table 4). One isolate was most closely related to a canine isolate from the UK. Three STs (ST38, ST10 and ST58) were shared by our study isolates and diarrhoeal isolate from the GEMS study (Supplementary Figure 2), with just eight alleles separating our commensal ST38 strain from a diarrhoeal isolate from the GEMS study (Figure 5).

**Table 4:** Closest relatives to the study isolates

| Sample ID | 7-gene ST | Neighbour host | Neighbour status | Neighbour's country of isolation | Allelic distance |
| --- | --- | --- | --- | --- | --- |
| H-32_5 | 10 | Human | Unknown | UK | 18 |
| H-36_1 | 59 | Human | Unknown | UK | 18 |
| H-39_1 | 452 | Human | Commensal | UK | 26 |
| H-9_1 | 2705 | Livestock |  | China | 29 |
| H-18_3 | 9274 | Human | Commensal | Unknown | 34 |
| H-2_1 | 9274 | Human | Commensal | Unknown | 34 |
| H-22_1 | 316 | Human | Commensal | UK | 35 |
| H-38_1 | 394 | Human | Pathogen (cystitis) | US | 39 |
| H-25_4 | 337 | Human | Unknown | Mali | 43 |
| H-37_1 | 5148 | Human | Pathogen (diarrhoea) | Ecuador | 43 |
| H-26_1 | 641 | Livestock |  | US | 46 |
| H-26_5 | 398 | Poultry |  | Kenya | 47 |
| H-48_2 | 485 | Human | Commensal | Tanzania | 57 |
| H-15_1 | 9277 | Human | Commensal | Zambia | 68 |
| H-15_2 | 747 | Human | Commensal | Egypt | 72 |
| H-28_1 | 469 | Human | Commensal | Kenya | 77 |
| H-21_2 | 1204 | Avian |  | Kenya | 81 |
| H-34_2 | 10 | Livestock |  | UK | 83 |
| H-38_2 | 58 | Human | Pathogen (bloodstream infection) | Australia | 87 |
| H-34_4 | 1727 | Unknown | Unknown | Unknown | 89 |
| H-35_1 | 226 | Human | Commensal | China | 93 |
| H-21_1 | 58 | Unknown | Unknown | Unknown | 98 |
| H-21_4 | 540 | Human | Unknown | Belgium | 100 |
| H-32_2 | 2175 | Livestock |  | UK | 100 |
| H-26_2 | 10 | Livestock |  | US | 111 |
| H-32_1 | 101 | Unknown | Unknown | Unknown | 111 |
| H-50_2 | 515 | Environment |  | Canada | 117 |
| H-41_1 | 43 | Unknown | Unknown | Unknown | 120 |
| H-26_4 | 2741 | Human | Commensal | Germany | 126 |
| H-50_1 | 410 | Livestock |  | US | 140 |
| H-18_1 | 38 | Poultry |  | US | 144 |
| H-21_5 | 223 | Unknown | Unknown | Unknown | 145 |
| H-40_1 | 155 | Unknown | Unknown | US | 146 |
| H-41_2 | 9283 | Environment | Commensal | US | 191 |
| H-36_4 | 9278 | Avian |  | Kenya | 208 |
| H-9_3 | 2914 | Canine |  | UK | 272 |
| H-9_5 | 29 | Unknown | Unknown | Unknown | 288 |
| H-34_1 | 603 | Laboratory |  | UK | 325 |
| H-55_1 | 9279 | Environment |  | Unknown | 333 |
| H-18_2 | 9281 | Unknown | Unknown | France | 430 |
| H-25_1 | 181 | Human | Commensal | Tanzania | 607 |

**Figure 3.**
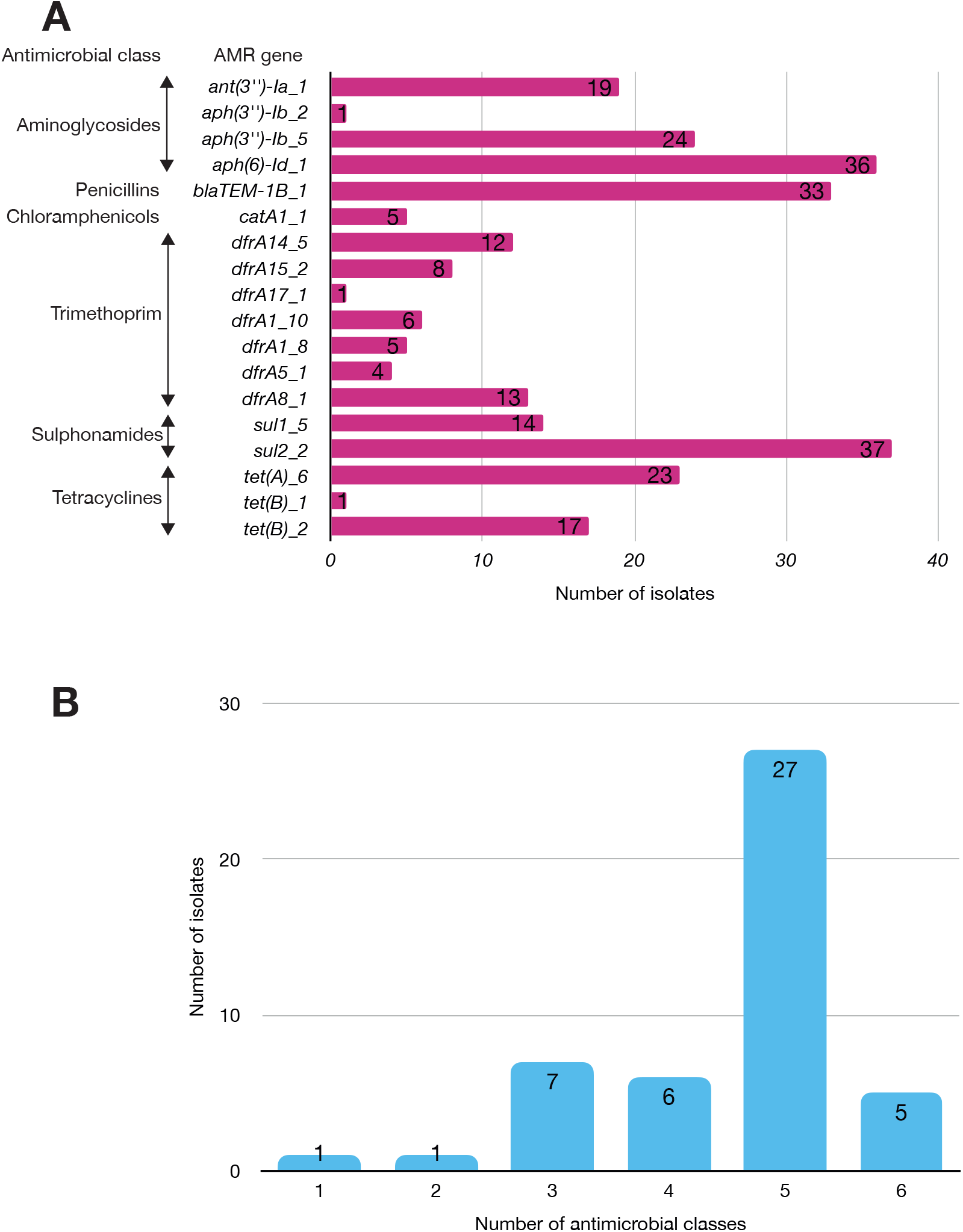
A: The prevalence of antimicrobial-associated genes detected in the isolates. The y-axis shows the detected AMR-associated genes in the genomes, grouped by antimicrobial class. B: A histogram depicting the number of antimicrobial classes to which resistance genes were detected in the corresponding strains.

**Figure 4.**
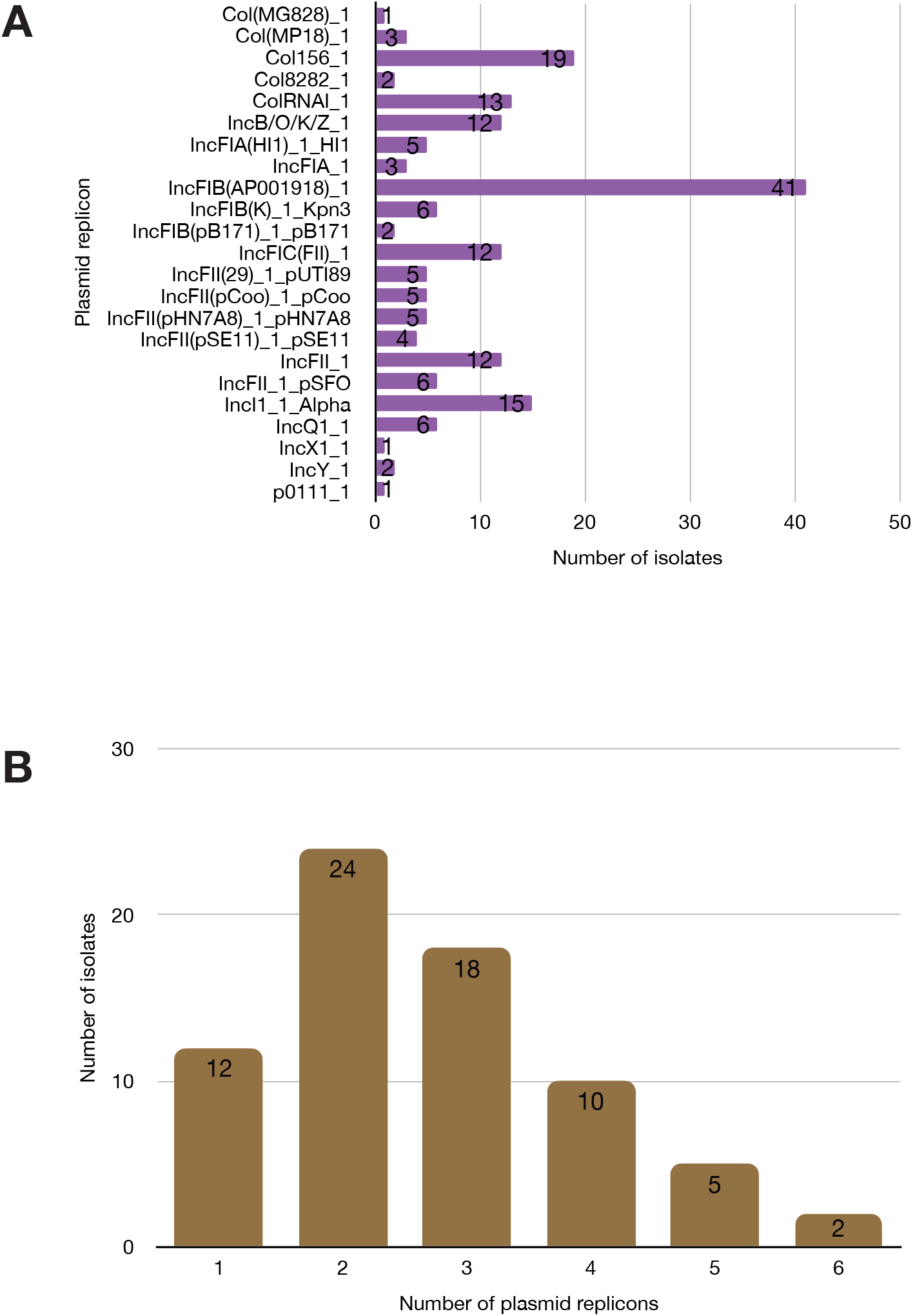
A: Plasmid replicons detected in the study isolates. B: A histogram depicting the number of plasmids co-harboured in a single strain.

**Figure 5.**
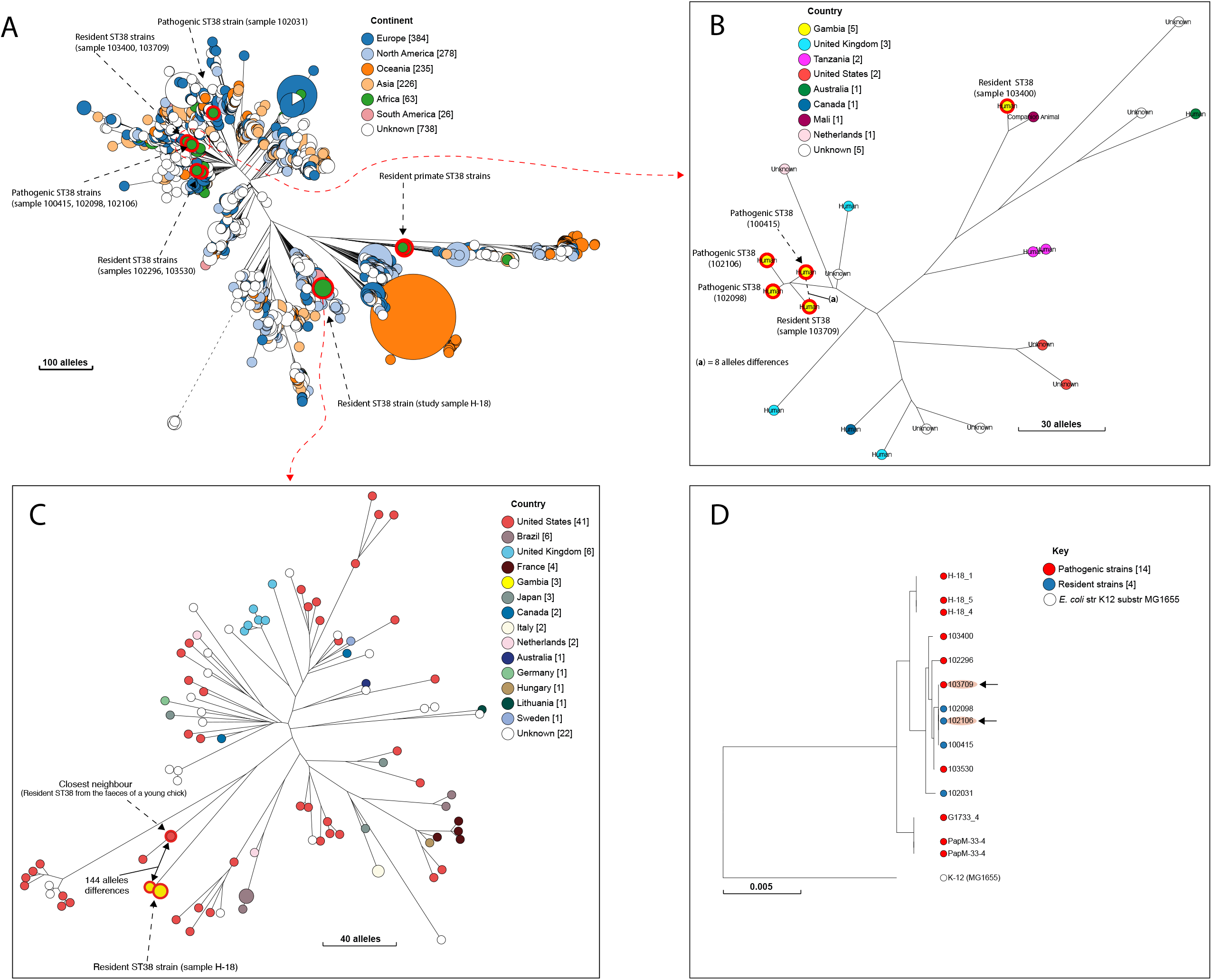
A: A NINJA neighbour-joining tree showing the population structure of *E. coli* ST38, drawn using the genomes found in the core-genome MLST hierarchical cluster at HC1100, which corresponds to ST38 clonal complex. B: The closest neighbour to a pathogenic strain reported in GEMS ^4^ is shown to be a commensal isolate recovered from a healthy individual. C: The closest relatives to the commensal ST38 strain recovered from this study is shown (red highlights), with the number of core-genome MLST alleles separating the two genomes displayed. D: A maximum-likelihood phylogenetic tree reconstructed using the genomes found in the cluster in C above, comprising both pathogenic and commensal ST38 strains is presented, depicting the genetic relationship between strain 100415 (pathogenic) and 103709 (commensal) (red highlights). The nodes are coloured to depict the status of the strains as pathogenic (red) or commensal (blue). The geographical locations where isolates were recovered are displayed in Figures 4A-C; the genome counts shown in square brackets.

We detected 130 genes encoding putative virulence factors across the 88 study isolates (Figure 2; Supplementary File 3). More than half of the isolates encoded resistance to three or more clinically relevant classes of antibiotics (Figure 3; Supplementary Figure 1). The most common resistance gene network was *-aph(6)-Id_1-sul2* (41% of the isolates), followed by *aph(3”)-Ib_5-sul2* (27%) and *bla-TEM-aph(3”)-Ib_5* (24%). Most isolates (67%) harboured two or more plasmid types (Figure 4). Of the 24 plasmid types detected, IncFIB was the most common (41%), followed by col156 (19%) and IncI_1-Alpha (15%). Nearly three-quarters of the multi-drug resistant isolates carried IncFIB (AP001918) plasmids, suggesting that these large plasmids disseminate resistance genes within our study population.

## Discussion

This study provides an overview of the within-host genomic diversity of *E. coli* in healthy children from a rural setting in the Gambia, West Africa. Surprisingly, we recovered a low rate of colonisation than reported elsewhere among children of similar age groups (42), with only a third of our study samples yielding growth of *E. coli*. This may reflect geographical variation but might also be some hard-to-identify effect of the way the samples were handled, even though they were kept frozen and thawed only just before culture.

Several studies have shown that sampling a single colony is insufficient to capture *E. coli* strain diversity in stools (20, 21, 23). Lidin-Janson *et al*. (43) claim that sampling five colonies provides a >99% chance of recovering dominant genotypes from single stool specimens, while Schlager *et al*. (24) calculate that sampling twenty-eight colonies provides a >90% chance of recovering minor genotypes. Our results confirm the importance of multiple-colony picks in faecal surveillance studies, as over half (57%) of our strains would have been missed by picking a single colony.

Although our strains encompassed all eight major phylotypes of *E. coli*, the majority fell into the A and B1 phylogenetic groups, in line with previous reports that these phylogroups dominate in stools from people in low- and middle-income countries (44, 45). The prevalence of putative virulence genes in most of our isolates highlights the pathogenic potential of commensal intestinal strains—regardless of their phylogroup—should they gain access to the appropriate tissues, for example, the urinary tract. Our results complement previous studies reporting genomic similarities between faecal *E. coli* isolates and those recovered from urinary tract infection (25, 46).

We found that within-host evolution plays a minor role in the generation of diversity, in line with Dixit et al. (20), who reported that 83% of diversity originates from immigration events, and with epidemiological data suggesting that the recurrent immigration events account for the high faecal diversity of *E. coli* in the tropics (47). Co-colonising variants belonging to the same ST tended to share an identical virulence, AMR and plasmid profile, signalling similarities in their accessory gene content. The estimated mutation rate for *E. coli* lineages is around one SNP per genome per year (48), so that two genomes with a most recent common ancestor in the last five years would be expected to be around ten SNPs apart. However, in two subjects, pairwise distances between genomes from the same ST (ST59 and ST5148) were large enough (14 and 18 respectively) to suggest that they might have arisen from independent immigration events, as insufficient time had elapsed in the child’s life for such divergence to occur within the host. However, it remains possible that the mutation rate was higher than expected in these lineages, although we found no evidence of damage to DNA repair genes. More than half of our isolates encode resistance to three or more classes of antimicrobials echoing the high rate of MDR (65%; confirmed by phenotypic testing) in the GEMS study. IncFIB (AP001918) was the most common plasmid Inc type from our study, in line with the observation that IncF plasmids are frequently associated with the dissemination of resistance (49). However, a limitation of our study is that we did not perform phenotypic antimicrobial resistance testing, although Doyle et al. (50) reported that only a small proportion of genotypic AMR predictions are discordant with phenotypic results.

Comparative analyses confirm the heterogeneous origins of the strains reported here, documenting links to other human commensal strains or isolates sourced from livestock or the environment. This is not surprising, as almost all study participants reported that animals are kept in their homes and children in rural Gambia are often left to play on the ground, close to domestic animals such as pets and poultry (51).

Our results show that the commensal *E. coli* population in the gut of healthy children in rural Gambia is richly diverse, with the independent immigration and establishment of strains contributing to the bulk of the observed diversity. Besides, this work has added significantly to the number of commensal *E. coli* genomes, which are underrepresented in public repositories. Although solely observational, our study paves the way for future studies aimed at a mechanistic understanding of the factors driving the diversification of *E. coli* in the human gut and what it takes to make a strain of *E. coli* successful in this habitat.

## Acknowledgements

We gratefully acknowledge the study participants in GEMS and all clinicians, field workers and the laboratory staff of the Medical Research Council Unit The Gambia at London School of Hygiene and Tropical Medicine involved in the collection and storage of stools in the GEMS study in Basse Field Station and Fajara.

## Data summary

All genomic assemblies for the strains included in this study are freely available from EnteroBase (http://enterobase.warwick.ac.uk/species/index/ecoli). The EnteroBase genome assembly barcodes are provided in Supplementary Files 1 and 2.

The raw genomic sequences have been deposited in the NCBI SRA, under the BioProject ID PRJNA658685 and accession numbers SAMN15880286 to SAMN15880281.

## Author contributions

Conceptualization: MA, MP; data curation, MP, NFA; formal analysis: EFN; analytical support: MD; funding: MA and MP; sample collection and storage: MJH, UNI, PET, CO; data management: SG; laboratory experiments, EFN, supervision, NFA, MP, JO, MA; manuscript preparation – original draft, EFN; review and editing, NFA, MP; review of the final manuscript, all authors.

## Supplementary material

**Supplementary Figure 1**

A co-occurrence matrix of acquired antimicrobial resistance genes detected in the study isolates. The diagonal values show how many isolates each individual gene was found in, while the intersections between the columns represent the number of isolates in which the corresponding antimicrobial resistance genes co-occurred.

**Supplementary Figure 2**

A Neighbour-joining phylogenetic tree depicting the genetic relationships among twenty-four strains isolated from diarrhoeal cases in the GEMS study ^4^. The Sequence types identified in these isolates are shown in the legend, with the genome count displayed in square brackets next to the respective sequence types. Three STs (ST38, ST58 and ST10) overlapped with what was found among commensal strains from this study (see Figure 2).

**Supplementary File 1**

Sequencing statistics and characteristics of twenty-four previously sequenced GEMS cases included in this study.

**Supplementary File 2**

A summary of the sequencing statistics of the study isolates reported in this study.

**Supplementary File 3**

A summary of the virulence factors detected among the study isolates and their known functions.

**Supplementary File 4**

A pairwise single nucleotide polymorphism matrix showing the SNP distances between the study genomes.

**Supplementary File 5**

List of the sample clones for which two independent cultures were obtained and sequenced, to find the SNPs between the same clones.

**Supplementary File 6**

Mutations in variants inferred to have been derived from within-host evolution.

