## Supplementary figures and images for "Genomic diversity of *Escherichia coli* isolates from healthy children in rural Gambia"

### Supplementary Figure 1

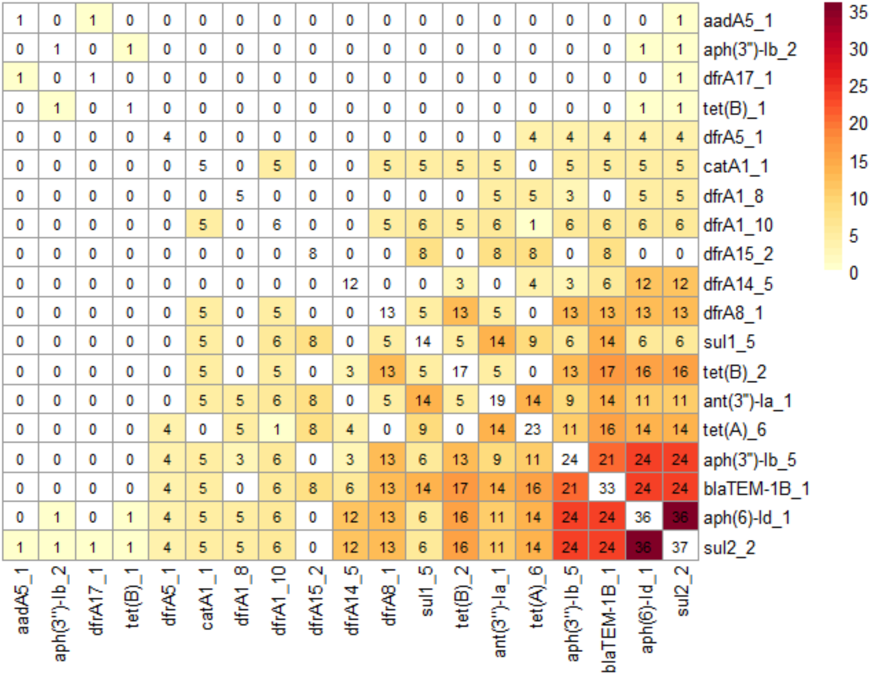

### Supplementary Figure 2

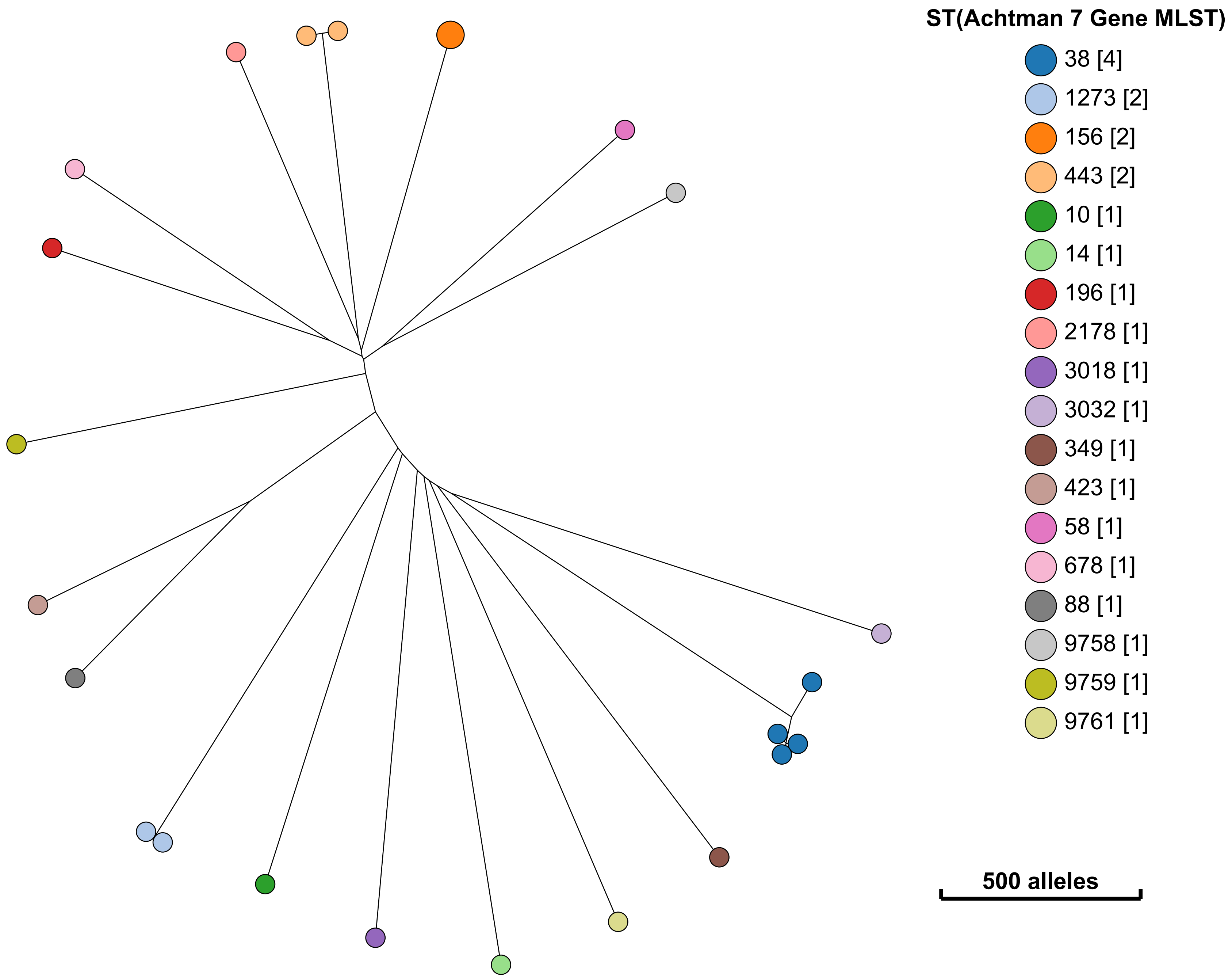
